# A Hypoxia Sensitive to Hypoxia Resistant Transformation in Long-Lived Germline Mutants

**DOI:** 10.1101/2021.06.22.449397

**Authors:** C Hemphill, E Pylarinou-Sinclair, O Itani, B Scott, MC Crowder, MR Van Gilst

## Abstract

Signals from the germline play a significant role in determining longevity in numerous animal models. In *C. elegans*, ablation of the germline leads to long life span and various other types of stress resistance. It has been reported that mutations that block oogenesis or an upstream step in germline development confer strong resistance to hypoxia. We report here that the hypoxia resistance of sterile mutants is dependent on developmental stage and age. In just a 12-hour period, sterile animals transform from hypoxia sensitive L4 larvae into highly hypoxia resistant adults. Since this transformation occurs in animals with no germline, the physiological programs that determine hypoxia sensitivity must occur independently of germline signals and instead rely on developmental signals from somatic tissues. Furthermore, we found two distinct mechanisms of hypoxia resistance in long-lived germline deficient animals. First, a DAF-16/FoxO *independent* mechanism that occurs in all hypoxia resistant sterile adults and, second, a DAF-16/FoxO *dependent* mechanism that confers an added layer of resistance, or “super-resistance”, to animals with no germline as they age past day 1 of adulthood. RNAseq data showed that nearly all genes involved in both cytosolic and mitochondrial protein translation, as well as in mitochondrial protein import, are repressed in germline deficient adults and further repressed as they age. The hypoxia super-resistance of aging germline deficient animals was suppressed by dual mutation of *ncl-1* and *larp-1*, two regulators of nucleolar biology and protein translation, demonstrating that the hypoxia super-resistance mechanism involves reduced protein translation. These studies provide novel insight into a profound physiological transformation that takes place in germline mutants during development, showing that some of the unique physiological properties of these long-lived animals are dependent on developmental repression of genes involved in protein translation, which operate independently of germline signals.

**AUTHOR SUMMARY:** In addition to being extremely long lived, germline deficient animals have other extraordinary properties, such as robust resistance to oxygen deprivation. Here we provide new insight into the mechanisms of hypoxia resistance in germline deficient animals. We demonstrate that, in just a 12-hour period, germline mutants transform from hypoxia sensitive larvae into highly hypoxia resistant adults. Therefore, hypoxia resistance is not a general property of germline ablated animals, but is instead “switched on” only in adult animals. We have found two distinct mechanisms of hypoxia resistance in germline deficient animals and both mechanisms are mediated by signals from somatic tissues and do not require the germline. We have determined that reduced transcription of genes involved in protein translation is one of the mechanisms of hypoxia resistance. Like hypoxia resistance, repression of protein translation genes only occurs in adults. Our findings establish that the unique physiological properties of germline-deficient animals are “switched” on in adults and therefore must be mediated by developmental signals from somatic tissues. We conclude that the L4/adult developmental switch in germline ablated animals presents an excellent system for investigating the longevity and hypoxia resistance of germline deficient animals.

## INTRODUCTION

Oxygen is critical for the function of nearly all living organisms. Accordingly, depriving an organism of oxygen can damage or kill cells and tissues and eventually lead to organismal death. In humans, cardiovascular events such as stroke and heart attack impede blood from reaching critical tissues resulting in ischemia and hypoxic injury. It is for this reason that hypoxic injury is intensively studied in a wide range of experimental models, including vertebrate animals, cell culture systems, as well as genetic model organisms such as *C. elegans* and *Drosophila*.

The unique advantage of *C. elegans* as an experimental model is its versatile genetic screening capabilities (Brenner 1974). Well-conceived random mutagenesis or RNAi screens can identify new genes involved in just about any physiological process. Like other animals, depriving *C. elegans* of oxygen leads to tissue damage and organismal death, thus nematodes can be an excellent model for studying hypoxic injury (Anderson, Mao et al. 2009). RNAi and mutagenic screens in *C. elegans* have identified over 200 genes that, when inhibited, enable animals to better survive oxygen deprivation (Anderson, Mao et al. 2009, Mabon, Mao et al. 2009, Itani, Zhong et al. 2021). The mechanisms by which most of these genes influence hypoxic injury have yet to be determined.

Perhaps the most well-studied pathways influencing hypoxic injury in *C. elegans* are protein translation and insulin signaling (Anderson, Mao et al. 2009, Mabon, Scott et al. 2009, Itani, Zhong et al. 2021). The role of protein translation appears to be the most critical, as mutagenesis screens designed to identify genes involved in hypoxic injury isolated multiple genes involved in protein synthesis, far more than any other gene class (Itani, Zhong et al. 2021). Reduced function mutations have been isolated in multiple tRNA synthetase genes, as well as other regulators of protein synthesis such as the RNA helicase gene *ddx-52*, a tRNA nuclear exporter gene *xpo-3*, a tRNA ligase gene *rtcb-1*, and an oligopeptide transporter gene *pept-1*. It has been confirmed that a hypoxia resistant loss of function mutation in the threonine tRNA synthetase *tars-1* leads to a reduction in the rate of protein synthesis (Itani, Zhong et al. 2021). A suppressor screen has isolated genes that can restore translation rates in these reduced translation mutants and has thereby expanded the understanding of regulatory networks that control protein translation and hypoxic injury (Itani, Zhong et al. 2021). Specifically, dual mutations in *larp-1* and *ncl-1*, regulators of protein translation, are able to suppress the hypoxia resistance of several protein translation mutants. Proteomic studies have shown that mutation of *ncl-1* and *larp-1* leads to an increase in ribosomal proteins which enables translation to be restored to wild type levels in tRNA synthetase mutants (Itani, Zhong et al. 2021). In summary, the study of protein translation and its link to hypoxia resistance in *C. elegans* has yielded novel insight into both hypoxic injury and protein translation.

Insulin signaling also has a significant impact on hypoxic injury. Specifically, loss of function mutations in the *daf-2* insulin receptor lead to strong hypoxia resistance (Mabon, Scott et al. 2009). This occurs through the standard insulin signaling pathway as a loss of function mutation in the FoxO gene homolog *daf-16* is able to suppress the hypoxia resistance of *daf-2* mutants. Mutation of *daf-16* does not suppress the hypoxia resistance of translation mutants, however, arguing that these two pathways influence hypoxia resistance through different mechanisms. Loss of function mutations in *daf-2* also lead to a prolonged lifespan, suggesting there may be a connection between the mechanisms of longevity and hypoxia resistance (Dorman, Albinder et al. 1995). However, mutations in protein translation genes that lead to hypoxia resistance have little impact on longevity, implying that hypoxia resistance doesn’t necessarily bring about increases in lifespan (Anderson, Mao et al. 2009).

Another factor that has a significant impact on hypoxic injury in *C. elegans* is reproduction (Mendenhall, LeBlanc et al. 2009). Specifically, any sterile mutation that prevents oogenesis or an upstream step in germline development confers strong resistance to hypoxia (Mendenhall, LeBlanc et al. 2009). For *glp-1(ts)* animals, which do not contain a germline when grown at 25°C, RNAi of AMPK subunits is able to partially suppress hypoxia resistance, potentially linking carbohydrate metabolism to the mechanism of sterile mutant hypoxia resistance (LaRue and Padilla 2011). Like *daf-2* mutants, germline ablated animals are long-lived raising the possibility that longevity and hypoxia resistance are related. However, mutation of *daf-16* suppresses the longevity of germline ablated animals whereas it has been reported that mutation of *daf-16* does not suppress the hypoxia resistance of *glp-1(ts)* mutants (Mendenhall, LeBlanc et al. 2009). Furthermore, other sterile mutants that have not been reported to be long-lived are resistant to hypoxia in a *daf-16* independent fashion, suggesting that longevity and hypoxia resistance are occurring through distinct mechanisms in sterile mutants. The link between sterility and hypoxia resistance is also an important one given that many of the 200+ hypoxia associated genes also partially reduce reproductive output when knocked down (Mabon, Mao et al. 2009). Therefore, knockdown of these genes may yield hypoxia resistance simply because of their impact on reproduction. It is not yet known, however, if partial sterility leads to partial hypoxia resistance, so it is unclear if these genes influence hypoxia by reducing reproductive output.

There is another connection between insulin signaling, germline development, and protein translation. Several long-lived mutants, including *daf-2(lf)* and *glp-1(ts)* mutants, possess smaller nucleoli (Tiku, Jain et al. 2017). In fact, smaller nucleoli, which play a role in the regulation and maturation of ribosomal RNAs, are a hallmark of longevity in multiple species. Consistent with the role of nucleoli in ribosome assembly and protein translation, fewer ribosomal proteins are present in both *daf-2* and *glp-1(ts)* mutants when compared to WT worms (Tiku, Jain et al. 2017). This observation suggests that it is likely that protein translation is reduced in both of these mutants, as well as other long-lived backgrounds. Even though reduced protein translation alone is not sufficient to prolong lifespan, numerous studies have shown that, in some cases, reduced protein translation leads to a lengthened lifespan (Hansen, Taubert et al. 2007, Pan, Palter et al. 2007, Syntichaki, Troulinaki et al. 2007). Mutation of *daf-16* is sufficient to suppress both nucleolar size and longevity in both *daf-2* and *glp-1(ts)* mutants. Smaller nucleoli in both mutants were also suppressed by mutation of *ncl-1,* which influences nucleolar physiology by repressing production of *fib-1*, a regulator of ribosomal RNA maturation (Tiku, Jain et al. 2017). Taken together, there are still many open questions regarding the relationships of reproduction and protein translation to hypoxic injury and longevity. Accordingly, we set out to further establish the mechanisms of hypoxia resistance in sterile mutants.

## RESULTS

### Germline Mutants are Resistant to Hypoxic Injury

A previous study has shown that sterile mutations that block oogenesis or an upstream step in germline development render animals resistant to hypoxic injury at 20°C (Mendenhall, LeBlanc et al. 2009). We set out to identify the mechanisms by which sterility leads to hypoxia resistance in *C. elegans.* We selected several different types of sterile strains to determine if mutations that prevent oogenesis also lead to hypoxia resistance under our conditions (because hypoxic injury is more robust and consistent at higher temperatures, we expose our worms to hypoxia at 26°C as opposed to 20°C as was previously reported for sterile mutants). The first type of sterile mutations tested included two temperature sensitive mutations, *glp-1(e2141)* and *glp-4(bn2)*. Worms containing either of these two mutations do not develop a germline when grown at or above 25°C. The second type of mutation was a temperature sensitive allele of the *fem-3* gene, *fem-3(q20)*, which leads to a masculinized germline that makes sperm but not oocytes when grown at or above 25°C. The third type of mutation was a different temperature sensitive allele of the *fem-3* gene, *fem-3(e2006)*, which leads to a feminized germline that makes oocytes but not sperm at or above 25°C. The fourth type of mutant was a *gld-1(lf)* mutant that blocks oogenesis by inhibiting stem cell differentiation leading to a tumorous germline full of undifferentiated mitotic cells. Strains containing all of these sterile mutations were tested for hypoxia sensitivity at 26°C and compared to wild type (WT) hermaphrodites, which produce both sperm and oocytes under our experimental conditions.

For our hypoxia assay, day 1 adults (12-24 hours after the L4/adult transition) were exposed to hypoxia for 24 or 32 hours, rescued on food containing plates and then scored for survival. Under these conditions, only about 6% of WT adults survive 24 hours of hypoxia and less than 1% survive 32 hours of hypoxia (Fig. 1A). Both the *glp-1(e2141)* and *glp-4(bn2)* mutant strains, which do not contain a germline, were highly resistant to hypoxia with >99% of day 1 adults surviving 24 hours of hypoxia and over 70% of day 1 adults surviving 32 hours of hypoxia. We found that the *fem-3(q20)* mutant, which makes sperm but not oocytes, was also resistant to hypoxic injury with >99% of day 1 adults surviving 24 hours of hypoxia and ∼30% surviving as much as 32 hours of hypoxia. Finally, we also found that the *gld-1(lf)* tumor strain was resistant to hypoxia, with >99% surviving 24 hours of hypoxia and ∼30% surviving 32 hours of hypoxia. In contrast, the *fem-3(e2006)* mutant, which makes oocytes but not sperm, displayed WT levels of hypoxia sensitivity (Fig 1A). Therefore, we found that mutations that prevent oogenesis were also highly resistant to hypoxic injury under our conditions. It should be noted that although four of five strains exhibited strong hypoxia resistance, we found that the germline deficient *glp-1(e2141)* and *glp-4(bn2)* mutants were significantly more resistant to hypoxia than the *fem-3(e2006)* masculinized strain and the *gld-1(lf)* tumor strain, which was most evident at 32 hours of hypoxia (Fig 1A).

**Figure 1.**
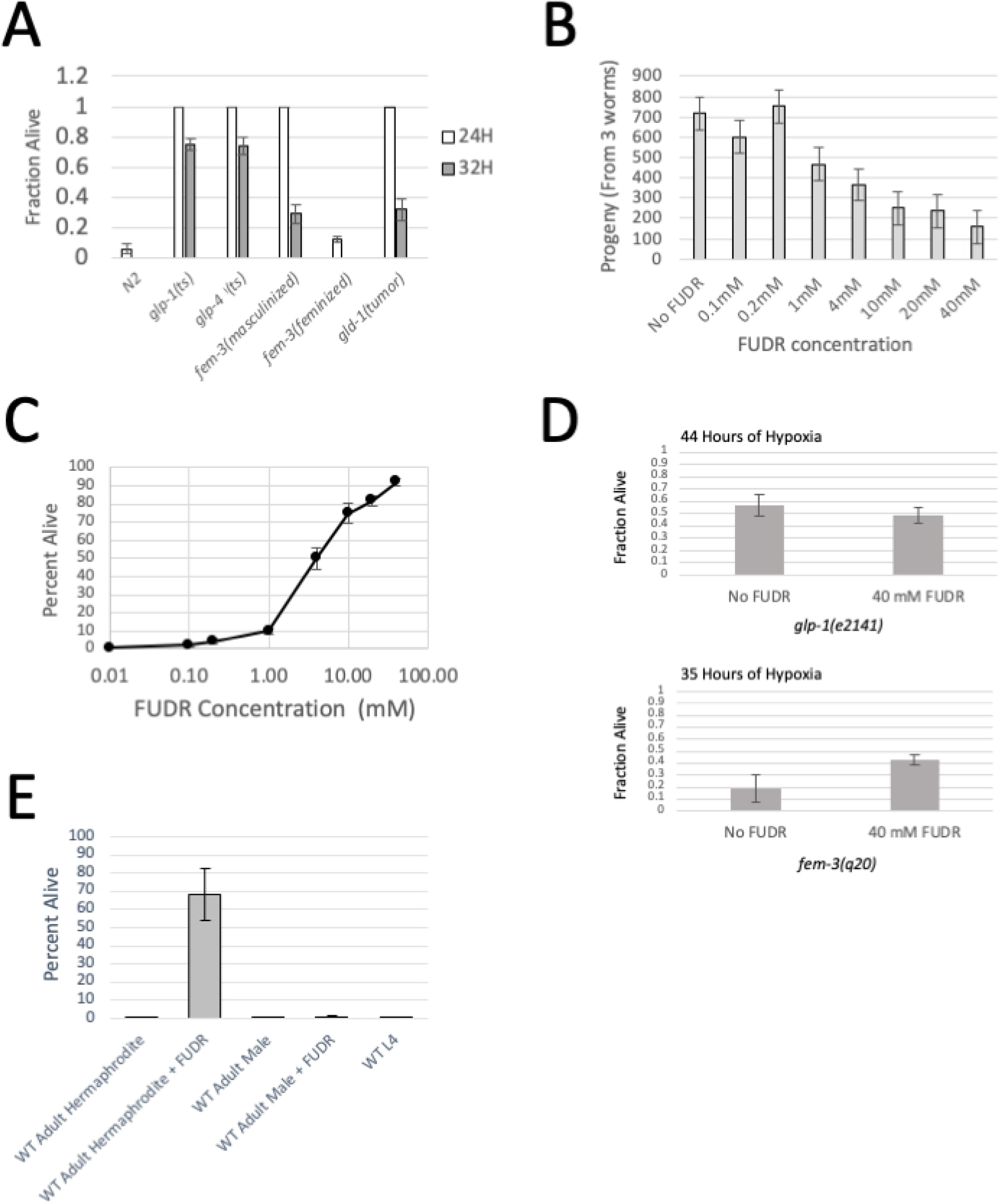
Hypoxia Resistance of Sterile Mutants. (A) The fraction of adult worms surviving either 24 hours or 32 hours of hypoxia at 26.5°C. All animals were grown to adulthood at the non-permissive temperature (25°C) to induce sterility. Data show that germline ablated animals (*glp-1(ts)* and *glp-4(ts)*), masculinized *fem-3(q20)* mutants (*fem-3(masculinized)*) and *gld-1(tumor)* mutants are highly resistant to hypoxic injury, whereas the feminized *fem-3(e2006)* mutant (*fem-3(feminized)*) displays close to wild type hypoxia sensitivity. Error bars represent standard error. (B) The average number of progeny produced by 3 WT worms when treated with different concentrations of FUDR. Data show that the brood size of WT worms decreases as they are exposed to higher concentrations of FUDR. The fertility mid-point is near 4 mM FUDR. Error bars represent standard error. (C) The percentage of WT worms surviving 24 hours of hypoxia when treated with different concentrations of FUDR. Data show that as FUDR concentration increases, so does hypoxia resistance, mid-point is 4mM FUDR. Error bars represent standard error (D) Fraction of *glp-1(ts)* and *fem-3(q20)* adults surviving hypoxia with and without 40 mM FUDR. Data show that FUDR increases the survival of *fem-3(q20)* worms, but not the survival of *glp-1(e2141)* adults. 44 hours of hypoxia were used for *glp-1(e2141)* adults and 35 hours of hypoxia were used for *fem-3(q20)* adults. These time points were chosen so that survival was in a range where the impact of FUDR would be visible. Standard error is shown. (E) The percentage of WT adult hermaphrodites, WT adult males, and hermaphrodite L4 larvae surviving 24 hours of hypoxia with and without FUDR treatment. WT hermaphrodites are resistant to hypoxia when treated with 40 mM FUDR, whereas WT males +/- FUDR are not resistant to hypoxic injury. WT L4 larvae are also not resistant to hypoxic injury. WT L4 larvae were not treated with FUDR because FUDR is toxic to *C. elegans* larvae (Gruber, Ng et al. 2009). Error bars represent standard error.

### Partial Sterility Leads to Partial Hypoxia Resistance in WT Animals

There are two possible ways that blocking oogenesis could lead to hypoxia resistance. First, hypoxia resistance may only occur if worms are completely prevented from reaching the oogenic stage of development, in which case worms would have to be completely sterile in order to be hypoxia resistant. Alternatively, hypoxia resistance could be proportional to the amount that oogenesis is blocked, in which case a partial reduction in fertility would lead to partial hypoxia resistance. Sterility can be chemically induced in *C. elegans* with floxuridine (FUDR). Importantly, reproductive capacity can be titrated by treating worms with different concentrations of FUDR. Therefore, we used a titration of FUDR to induce partial sterility in wild type worms and to determine if partial sterility leads to hypoxia resistance. We found that reproductive output was not reduced at FUDR concentrations below 0.2 mM but was partially suppressed at concentrations ranging from 0.2 mM FUDR to 40 mM FUDR with a midpoint of ∼4 mM FUDR (Fig 1B). The sterility brought about by FUDR correlated well with the amount of hypoxia resistance. Specifically, we found that concentrations of 0.2 mM FUDR or less had little impact on hypoxia resistance, whereas concentrations between 0.2 mM FUDR and 40 mM FUDR had an increasing impact on hypoxia resistance with 4 mM FUDR lying at the midpoint of the curve (Fig 1C). Therefore, hypoxia resistance does not require complete sterility and partial sterility leads to partial hypoxia resistance.

If FUDR is impacting hypoxia resistance through the same mechanism as the sterile mutants, we would expect that FUDR would not further increase the hypoxia resistance of sterile mutants. Therefore, we exposed both *glp-1(e2141)* and *fem-3(q20)* to FUDR to determine if FUDR could increase the hypoxia resistance of sterile mutants. We found that FUDR did not make *glp-1(e2141)* worms more resistant to hypoxia, but did give a small but significant increase to the hypoxia resistance of *fem-3(q20)* mutants (Fig 1D). These results imply that inhibition of oogenesis alone is not sufficient to give full hypoxia resistance. Indeed, *fem-3(q20)* mutants still undergo germ cell proliferation and differentiation into sperm. Therefore, FUDR is still able to act on the partially differentiated germline of the *fem-3(q20)* mutants to bring about additional hypoxia resistance. In contrast, the sterility of *glp-1(e2141)* mutants is sufficient to get full hypoxia resistance. In this case, FUDR is unable to enhance hypoxia resistance in *glp-1(e2141)* animals because there are no germ cells for FUDR to act on.

### Oogenesis is Not Required for Hypoxia Sensitivity

Our results, consistent with previous studies, demonstrate that sterile mutations that prevent oogenesis in adult animals are sufficient to cause hypoxia resistance. This phenomenon suggests that oogenesis may be required for worms to be sensitive to hypoxia. If this hypothesis were true, we would expect that worms that do not make oocytes, such as hermaphrodite L4 larvae and WT males, should also be more resistant to hypoxic injury than WT adults. To address this issue, we subjected WT males and hermaphrodite L4 larvae to 24 hours of hypoxia and scored animals after 24 hours of recovery. We found that neither adult males nor hermaphrodite L4 larvae were more resistant than WT day 1 adults (Fig 1E). These results demonstrate that oogenesis is not a requirement for hypoxia sensitivity at 26°C.

We also found that growing WT males on 40 mM FUDR, a concentration that renders hermaphrodites resistant to hypoxia, was not sufficient to make WT males resistant to hypoxic injury (Fig 1E). Therefore, FUDR is only effective at making WT hermaphrodites resistant to hypoxia. Taken together, these results imply that it is the prevention of oogenesis in otherwise oogenic worms that leads to hypoxia resistance, not the lack of oogenesis *per se*.

### Sterility Dependent Hypoxia Resistance Only Occurs in Adults

When testing the hypoxia sensitivity of larvae and adults, we observed an interesting phenomenon for the sterile mutants. Specifically, we found that the *glp-1(e2141)* and *fem-3(q20)* mutant strains, which are both very resistant to hypoxia as day 1 adults, were not significantly resistant to hypoxia as L4 larvae (Fig 2A). In a 24-hour exposure to hypoxia, *glp-1(e2141)* and *fem-3(q20)* L4 larvae displayed WT sensitivity to hypoxia, with less than 10% of L4 larvae from both strains surviving 24 hours of hypoxia. In contrast, 100% of *glp-1(e2141)* day 1 adults and >95% of *fem-3(q20)* day 1 adults survived the same 24-hour exposure to hypoxia. The difference between L4 larvae and day 1 adults was even more apparent at longer hypoxia exposures. Indeed, less than 1% of the L4 larvae from both sterile mutants survived 32 hours of hypoxia, whereas ∼60% of *glp-1(e2141)* day 1 adults and ∼20% of *fem-3(q20)* day 1 adults survived the same 32-hour exposure (Fig 2A). Therefore, in just the time it takes to develop from L4 larvae to young adult (∼12 hours), the *glp-1(e2141)* and *fem-3(q20)* mutants transition from hypoxia sensitive L4 larvae to robustly hypoxia resistant day 1 adults. Such a transition is not observed in WT animals, however, as WT L4 larvae and day 1 adults displayed similar levels of hypoxia sensitivity after 24 hours of hypoxia exposure (Fig. 2A). We note that these results are further evidence that oogenesis is not required for hypoxia sensitivity, as *glp-1(e2141)* and *fem-3(q20)* L4 larvae do not undergo oogenesis, yet they still display wild-type sensitivity to hypoxic injury.

**Figure 2.**
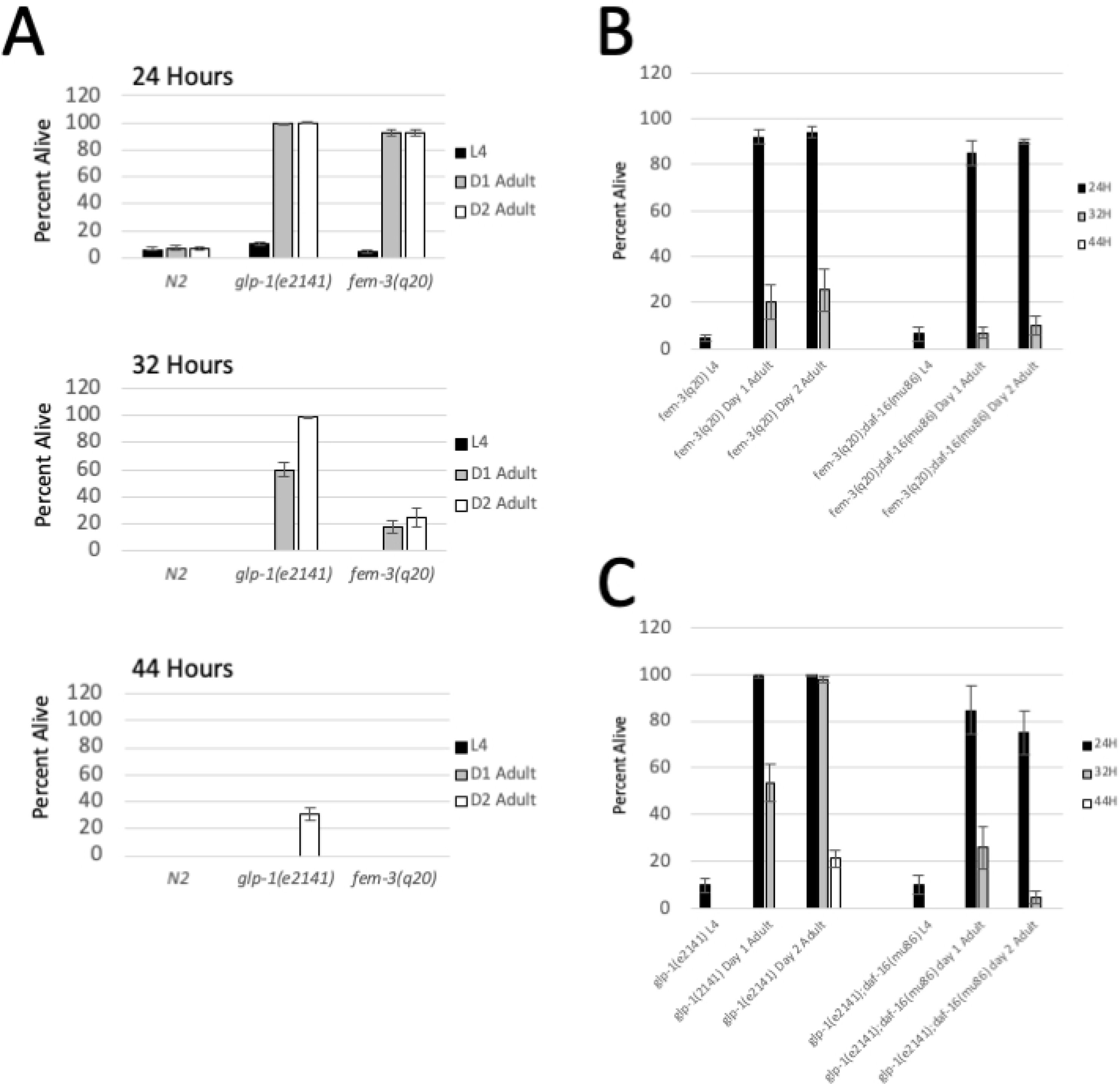
Two Mechanisms of Hypoxia Resistance in Aging Animals. (A) The percentage of worms surviving 24, 32, or 44 hours of hypoxia at 26.5°C. Worms from each genetic background were assayed as L4 larvae, day 1 adults, or day 2 adults. Germline ablated animals (*glp-1(e2141)*) become resistant to hypoxic injury as day 1 adults and even more resistant (super-resistant) as they age to day 2 of adulthood. In contrast, masculinized *fem-3(q20)* mutants become resistant to hypoxic injury as day 1 adults, but do not become super-resistant as they age to day 2 of adulthood. Error bars represents standard error. (B) Percentage of *fem-3(q20)* and *fem-3(q20);daf-16(mu86)* mutants surviving 24, 32, or 44 hours of hypoxia. The graph shows the survival of *fem-3* mutants as L4, day 1 adults, and day 2 adults (C) Percentage of *glp-1(e2141)* and *glp-1(e2141);daf-16(mu86)* mutants surviving 24, 32 or 44 hours of hypoxia. Data show that primary hypoxia resistance present in both *glp-1(e2141)* and *fem-3(q20)* mutants is not dependent on *daf-16*, however the aging-dependent super-resistance of day 3 *glp-1(e2141)* mutants is completely dependent on *daf-16*. Error bars represent standard error.

### *Glp-1(e2141)* Mutants Become “Super-Resistant” to Hypoxia as they Age

When testing the impact of developmental stage on hypoxia resistance, we also found that the hypoxia resistance of germline ablated animals increased significantly as adults aged to day 2 of adulthood (36-48 hours after the L4/adult transition). Indeed, >99% of *glp-1(e2141)* day 2 adults survived 32 hours of hypoxia compared to only ∼60% of day 1 adults (Fig 2A). Remarkably, ∼30% of day 2 adults survived as much as 44 hours of hypoxia, compared to less than 1% of *glp-1(e2141)* day 1 adults. Therefore, for the *glp-1(e2141)* mutant, L4 larvae display wild-type sensitivity to hypoxia, day 1 adults display strong resistance to hypoxia, and day 2 adults display even stronger resistance to hypoxia. We have called this aging dependent increase in hypoxia resistance “super-resistance”, as this is the strongest hypoxia resistance we have observed. Hypoxia resistance did not further increase when animals aged to day 3 of adulthood (data not shown).

The aging dependent super resistance phenomenon was limited to the *glp-1(e2141)* mutant, however, as *fem-3(q20*) mutants displayed a significant increase in hypoxia resistance between L4 and day 1 of adulthood, but saw no further increase in hypoxia resistance as they aged to day 2 of adulthood (Fig 2A). Taken together, these results suggest that there are multiple mechanisms of hypoxia resistance in sterile animals. A mechanism of primary hypoxia resistance which occurs in animals with inhibited oogenesis between L4 and day 1 of adulthood, and a second mechanism of “super-resistance” which we have only observed in germline deficient mutants as they age to day 2 of adulthood.

### Super Resistance of *glp-1(e2141)* Mutants is Dependent on Insulin Signaling

The insulin signaling pathway, operating through the DAF-16/FoxO transcription factor, is involved in a wide range of aging and stress related phenomena. Indeed, it is established that the longevity of *glp-1(e2141)* mutants is dependent upon DAF-16/FoxO (Hsin and Kenyon 1999). To determine if the hypoxia resistance of sterile mutants is also dependent upon *daf-16*, we acquired a *daf-16(mu86);glp-1(e2141)* double mutant strain and we built a *fem-3(q20);daf-16(mu86)* double mutant strain. We found that mutation of *daf-16* only mildly suppressed the hypoxia resistance of *glp-1(e2141)* and *fem-3(q20)* day 1 adults (Fig 2B and 2C). In contrast, we found that the aging dependent increase in hypoxia resistance in the *glp-1(e2141)* strain was completely suppressed by *daf-16*. Specifically, day 2 *glp-1(e2141)* adults were considerably more resistant to hypoxia than day 1 adults, but day 2 *daf-16(mu86);glp-1(e2141)* double mutants were no more resistant to hypoxic injury than day 1 adults (Fig 2C).

These results further solidify the notion that two distinct mechanisms of hypoxia resistance are operating in sterile mutants. First, a *daf-16 independent* mechanism that gives hypoxia resistance to both *fem-3(q20)* and *glp-1(e2141*) mutants after the transition from L4 larvae to day 1 of adulthood, this mechanism is consistent with what has previously been reported (Mendenhall, LeBlanc et al. 2009). Second, a *daf-16 dependent* mechanism that provides super-resistance to *glp-1(e2141)* mutants when they reach day 2 of adulthood. The *daf-16* dependent super resistance mechanism operates in the aging *glp-1(e2141)* mutant, but not in the masculinized *fem-3(q20)* mutant, and has not previously been reported.

### Germline Mutations Prevent Hypoxia Dependent Mitochondrial Protein Aggregation

Previous studies have shown that mitochondrial protein aggregation is one of the physiological hallmarks of hypoxic injury in *C. elegans* (Kaufman and Crowder 2015, Kaufman, Wu et al. 2017). Mitochondrial protein aggregation has been observed using transmission electron microscopy or by using a GFP reporter strain that expresses the mitochondrial UCR-11 protein (ubiquinol-cytochrome C reductase) fused to GFP in the body wall muscle. Mitochondrial protein aggregation is often accompanied by a swelling/rounding of body wall muscle mitochondria. Both mitochondrial protein aggregation and swelling are reversible under sublethal conditions, as mitochondria return to normal shape and aggregation is reversed 24 hours after return to normal oxygen conditions (Kaufman and Crowder 2015). Although previous studies have shown a correlation between mitochondrial protein aggregation and hypoxia sensitivity, it is not yet known if mitochondrial protein aggregates cause hypoxic injury and whether or not hypoxia resistant mutants prevent the formation of protein aggregates.

To examine mitochondrial protein aggregation in germline mutants, we crossed a *ucr-11::gfp* reporter strain into the *glp-1(e2141)* and *fem-3(q20)* mutant backgrounds. We then observed hypoxia dependent mitochondrial protein aggregation in both mutant reporter strains and compared them to the WT reporter strain. To account for the super-resistance mechanism, we examined both day 1 and day 2 adults of each strain. For WT animals, we found that 15 hours of hypoxia was sufficient to cause rounding/swelling of mitochondria along with the formation of abundant protein aggregates (Fig. 3A). Quantification of protein aggregates in WT animals revealed nearly 50 aggregates per high powered field (HPF) after 15 hours of hypoxia (Fig 3B). We observed no differences between day 1 and day 2 WT worms. In contrast, 15 hours of hypoxia led to no mitochondrial swelling or protein aggregation and less than 5 aggregates/HPF in the *fem-3(q20)* mutant and no swelling or detectable aggregates in the *glp-1(e2141)* background (Fig. 3A). For both germline mutants, longer periods of hypoxia exposure were required to cause significant mitochondrial protein aggregation. For *fem-3(q20)* animals and *glp-1(e2141)* day 1 adults, 24 hours of hypoxia were sufficient to cause mitochondrial swelling and significant protein aggregation, and for *glp-1(e2141)* day 2 adults, 30 hours of hypoxia were required to cause swelling and significant mitochondrial protein aggregation (Fig 3A and 3B). WT animals did not survive either a 24-hour or 30-hour hypoxia exposure. The fact that day 2 *glp-1(e2141)* animals are more resistant to mitochondrial protein aggregation than day 1 *glp-1(e2141)* animals implies that both the primary sterility dependent mechanism and the super-resistance mechanism protect against mitochondrial protein aggregation.

**Figure 3.**
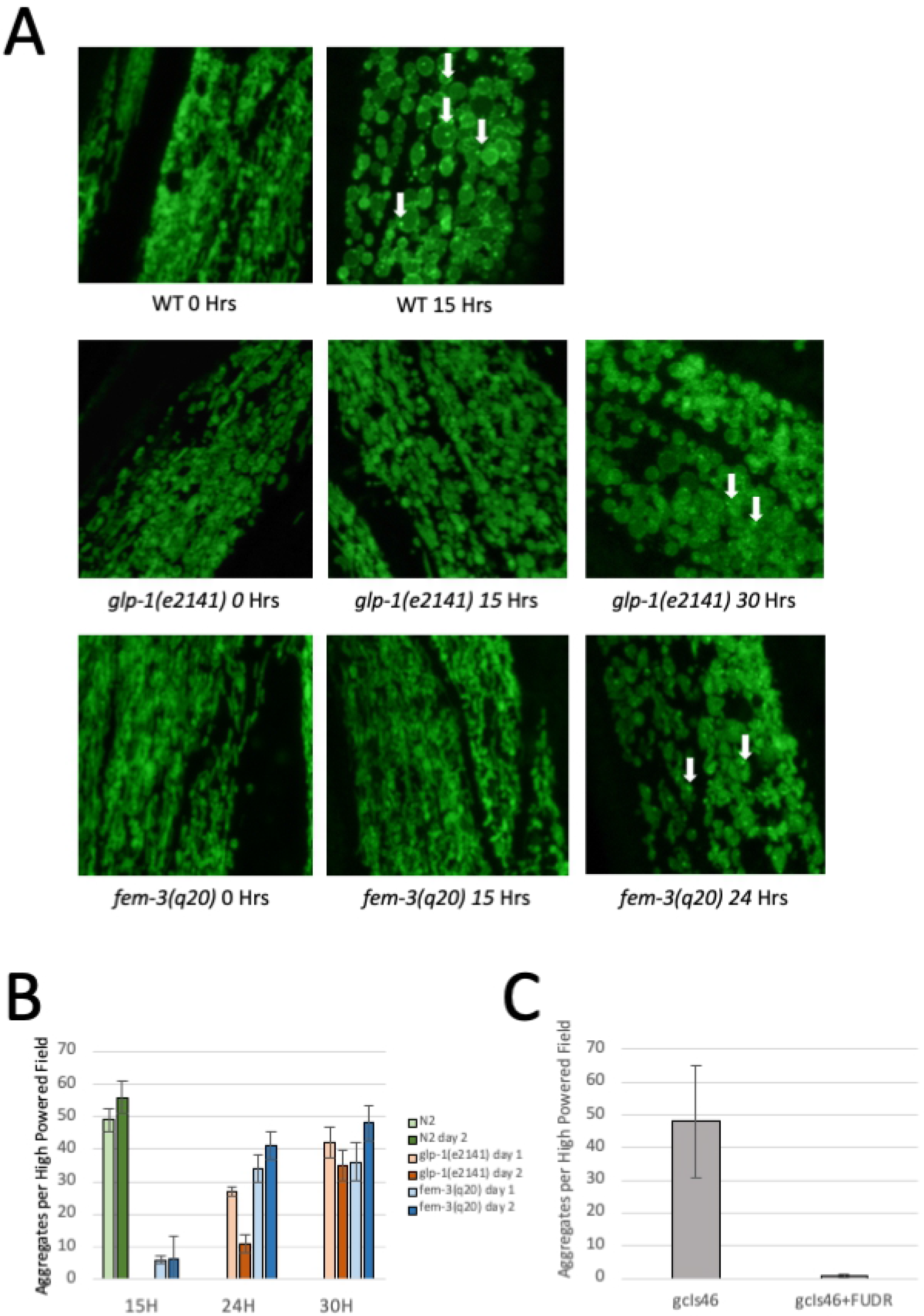
Sterile Mutants Prevent Hypoxia Dependent Mitochondrial Protein Aggregation. (A) GFP fluorescence images of WT, *glp-1(e2141)* and *fem-3(q20)* day 1 adults. Fluorescence is from a GFP tagged UCR-11 protein which localizes to body wall muscle mitochondria. The identity of the strain being shown, along with the time of hypoxia exposure is written below each image. Examples of mitochondrial protein aggregates are indicated by the white arrows. 15 hours of hypoxia is sufficient to cause rounding of the mitochondria and significant mitochondrial protein aggregation in WT worms, but not in *glp-1(e2141)* or *fem-3(q20)* mutants. 24 and 30 hours of hypoxia are sufficient to cause mitochondrial rounding and protein aggregation in *fem-3(q20)* and *glp-1(e2141)* adults respectively. (B) Quantification of protein aggregation after hypoxia. The average number of aggregates observed per high-powered field is displayed. The quantity of aggregates is shown after 15, 24, or 30 hours of hypoxia at 26.5°C. N/A is displayed for WT worms at 24 and 30 hours because WT worms do not survive these longer hypoxia exposures. Error bars represent standard error (C) Quantification of protein aggregates in WT worms after 15 hours of hypoxia at 26.5°C with and without FUDR. Data show that FUDR nearly eliminates mitochondrial protein aggregation in WT worms after hypoxia exposure. Error bars represent standard error.

We also used FUDR to determine if chemically induced sterility also prevented mitochondrial protein aggregation. WT animals carrying the *ucr-11::gfp* reporter were treated with 40 mM FUDR and then subjected to 15 hours of hypoxia. 15 hours of hypoxia were sufficient to induce significant protein aggregation in untreated animals, whereas animals treated with 40 mM FUDR displayed very little protein aggregation (Fig. 3C). Taken together, these results show that either sterile mutations or compounds that cause sterility are sufficient to prevent hypoxic injury and reduce the amount of hypoxia induced mitochondrial protein aggregation.

### RNAseq Analysis of Development and Aging in Germline Mutants

The most evident strategy for identifying changes in gene expression associated with hypoxia resistance would be to compare hypoxia sensitive WT animals to hypoxia resistant sterile mutants. There are significant disadvantages to making such comparisons, however, because WT worms possess an entire germline along with all of its associated gene expression, whereas *glp-1(e2141)* mutants possess no germline expression at all. Therefore, it is to be expected that there would be massive differences in gene expression solely due to the presence or absence of a germline, which would make it exceedingly difficult to isolate gene expression changes specifically associated with hypoxia resistance. Our findings gave us a much more attractive alternative for gene expression analysis. Indeed, in just a 12-hour period, *glp-1(e2141)* mutants transition from hypoxia sensitive L4 larvae into highly resistant adults, and then in another 24 hours *glp-1(e2141)* mutants become super-resistant to hypoxia. Thus, gene expression can be compared in the same genetic background at three different time points, eliminating complications associated with gene expression in the germline. For these reasons, we carried out RNAseq analysis to examine changes in gene expression that occurred in *glp-1(e2141)* animals between the hypoxia sensitive L4 larval stage of development, the hypoxia resistant day 1 of adulthood, and the super hypoxia resistant day 3 of adulthood.

The most striking result of our RNAseq analysis was the regulation of genes involved in protein translation. In *glp-1(e2141)* mutants every single annotated ribosomal gene, including both the cytosolic and mitochondrial ribosomal subunit genes, was repressed by greater than 2-fold during the L4 to adult transition, and then further repressed by day 3 of adulthood (Supplementary Tables 1 and 2). Interestingly, we observed that the cytosolic ribosomal subunit genes and the mitochondrial ribosomal subunit genes were repressed to different degrees, with the mitochondrial genes being repressed more robustly than the cytosolic ribosomal genes. Indeed, when plotted as a histogram based on the degree of repression, the cytosolic ribosomal genes form a tight distribution with most genes centered between 2.5-fold and 3-fold repression (mean = 2.67-fold repression) (Fig 4A). In contrast, the mitochondrial ribosomal subunit genes form a distinct distribution centered between 3.5-fold and 4-fold repression (mean = 3.81-fold). Both of these distributions are significantly different than the distribution of total *C. elegans* genes, which is centered between 1.0-fold and 1.5-fold repression/activation (Fig 4A). RNAseq data shows that the repression of cytosolic and mitochondrial ribosomal subunit genes increased dramatically as animals aged to day 3 of adulthood (Fig. 4B). Comparing L4 worms to day 3 adults, cytosolic ribosomal subunit genes formed a tight distribution with most genes centered between 3-fold and 4.5-fold repression (mean = 3.97-fold) and the mitochondrial ribosomal genes formed a tight distribution with most genes falling between 5-fold and 8-fold repression (mean = 6.91-fold). Therefore, it is clear that all ribosomal and mitochondrial ribosomal protein genes are significantly repressed in *glp-1(e2141)* mutants as they develop from L4 to adulthood, and further repressed as they age past day 1 of adulthood.

**Figure 4.**
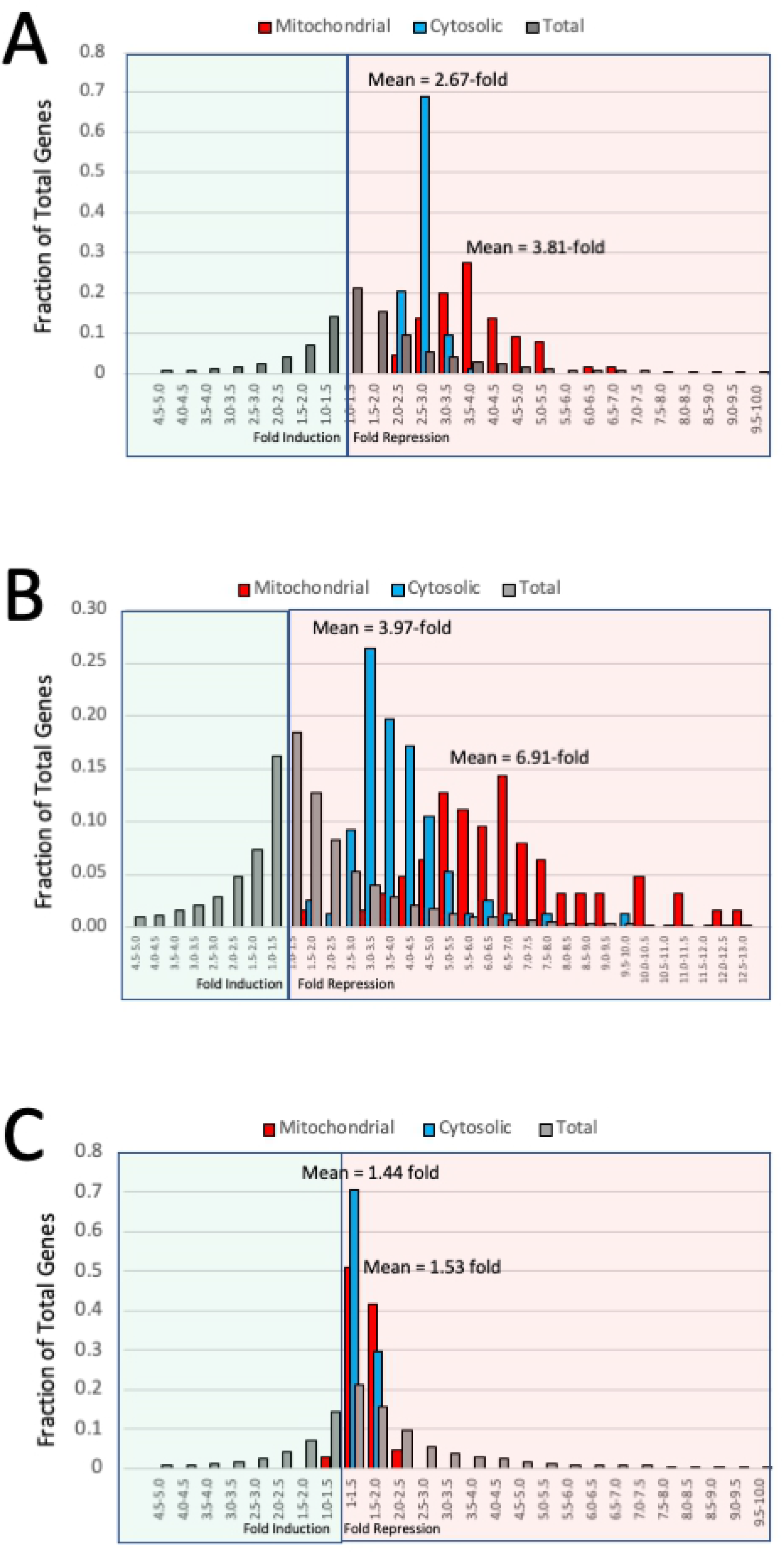
Ribosomal Gene Proteins (RPGs) are Repressed in *glp-1(e2141)* Adults. (A) Histogram of ribosomal genes binned by fold repression after the L4 to day 1 adult transition in *glp-1(e2141)* adults. Blue bars represent the fraction of the total ribosomal gene proteins (RPGs) in each range of gene repression or induction. Red bars represent the fraction of total mitochondrial ribosomal proteins (mitoRPGs) in each range of gene repression or induction. Gray bars represent the fraction of total genes in each range of repression or induction. Mean is the average repression of all RPGs or mitoRPGs. Data show that RPGs and mitoRPGs are repressed in two distinct groups with different mean repressions. (B) Histogram of ribosomal genes binned by fold repression between L4 *glp-1(e2141)* larvae versus day 3 adults. Repression between L4 larva and day 3 adults is considerably stronger than between L4 larva and day 1 adults, with the distributions of both RPGs and mitoRPGs shifting significantly higher (C). L4 *fem-3(q20)* vs day 1 *fem-3(q20)*. There are no RPGs or mitoRPGs repressed by more than 2-fold in *fem-3(q20)* animals between L4 larval stage and day 1 of adulthood. The distributions of RPGs and mitoRPGs in *fem-3(q20)* animals are not significantly different than the distribution of total genes.

### Multiple Steps in a Nucleolar Regulatory Pathway are Repressed in *glp-1(e2141)* Adults

In addition to the ribosomal subunit genes, other genes involved in protein translation were repressed in *glp-1(e2141)* adults. In fact, 27 out of 32 tRNA synthetase genes were repressed by more than 2-fold in day 1 adults (mean = 2.8-fold repression) (Supplemental Figure 1). Additionally, 11 ribosomal factors were repressed in day 1 *glp-1(ts)* adults. Interestingly, the tRNA synthetase genes and ribosomal factors were repressed in day 1 adults, but were not further repressed in day 3 adults, exhibiting a regulatory pattern different from that of the ribosomal and mitochondrial ribosomal genes. Other genes involved in translation included regulators of mitochondrial translation activation and elongation, which were repressed via the same pattern as the mitochondrial ribosomal genes, i.e., repressed in day 1 adults by ∼4-fold and further repressed by ∼8-fold in day 3 adults (Fig 5A and Table 1).

**Figure 5.**
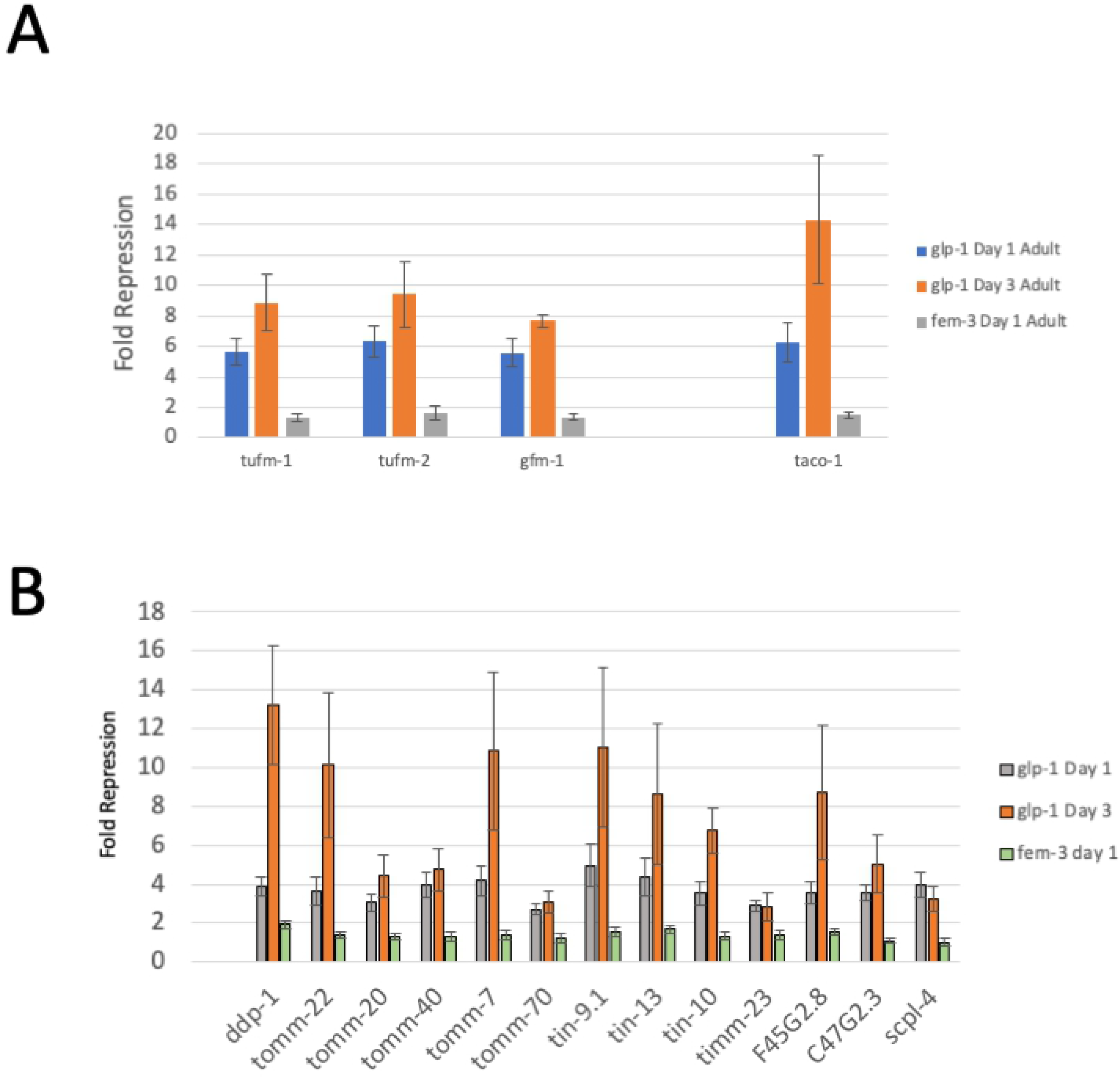
Other Translation Genes Regulated in *glp-1(e2141*) Mutants. (A) Repression of mitochondrial translational regulatory genes between the L4 larval stage and day 1 or day 3 of adulthood in *glp-1(e2141)* mutants or between the L4 larval stage and day 1 of adulthood in *fem-3(q20)* mutants. Error bars are standard errors. (B) Repression of mitochondrial translocase genes between the L4 larval stage and day 1 or day 3 of adulthood in *glp-1(e2141)* animals or between the L4 stage and day 1 of adulthood in *fem-3(q20)* mutants. Error bars represent standard errors.

**Table 1.**

|  | *glp-1(e2141)* |  | *fem-3(q20)* |
| --- | --- | --- | --- |
|  | L4 -> d1 | L4 -> d3 | L4 -> d1 |
| Cytosolic Ribosomal Genes | 2.7 +/- 0.25 | 4.0 +/- 1.24 | 1.4 +/- 0.21 |
| tRNA Synthetase Genes | 2.8 +/- 0.85 | 3.0 +/- 1.08 | 1.0 +/- 0.18 |
| Nucleolar Genes | 5.6 +/- 2.60 | 4.2 +/- 1.71 | 1.2 +/- 0.14 |
| Other Ribosomal Factors | 4.7 +/- 1.79 | 4.5 +/- 2.24 | 1.2 +/- 0.19 |
| Mitochondrial Ribosomal Genes | 3.8 +/- 0.95 | 6.9 +/- 2.44 | 1.5 +/- 0.30 |
| Mitochondrial Translation Factors | 6.0 +/- 0.40 | 10.0 +/- 2.9 | 1.5 +/- 0.13 |
| Mitochondrial Translocases | 3.7 +/- 0.17 | 7.1 +/- 0.97 | 1.4 +/- 0.07 |

Average fold repression of translation gene types between L4 and d1 or d3 of adulthood, standard deviation is shown. Three classes of gene regulation are observed, indicated by color.

We also identified eleven genes involved in nucleolar function, including two regulators of nucleolar function, the *fib-1* gene and the *let-7* microRNA, as repressed in day 1 adults (Supplemental Figure 1). Remarkably, *let-7* is repressed in d1 adults by more than 100-fold. *Let-7* has been reported to influence *fib-1* expression by binding to the *ncl-1* mRNA and repressing post transcriptional expression of the *ncl-1* gene (Gruber, Ng et al. 2009, Yi, Ma et al. 2015). The *ncl-1* gene product then represses expression of *fib-1*. Repression of *let-7* would be predicted to increase the production of the NCL-1 protein, which would then repress transcription of the *fib-1* gene. The fact that both the *let-7* microRNA and the *fib-1* gene are repressed in *glp-1(e2141)* adults fits with this model. Therefore, multiple components of this established nucleolar regulatory pathway are dramatically impacted by the development and aging of *glp-1(e2141)* animals.

Taken together, we have identified three classes of gene regulation for genes involved in protein translation (Table 1). First, the ribosomal genes, which are repressed ∼2.7-fold in day 1 adults and further repressed ∼4.0-fold in day 3 adults. Second, mitochondrial ribosomal genes and translation factors, which are repressed more strongly at ∼3.8-fold in day 1 adults and further repressed by ∼6.9-fold in day 3 adults. Third, tRNA synthetases, ribosomal factors, and nucleolar genes, which are repressed in day 1 adults, but *not further repressed* in day 3 adults.

### Mitochondrial Protein Translocation Genes are Co-regulated with Ribosomal Genes

Our results show that genes that function together in translation are regulated together in developing and aging *glp-1(e2141)* mutants. Thus, we asked if any other gene families were regulated in a similar fashion to the translation genes, as these genes may also be functionally related to translation. Indeed, we found that all of the genes involved in mitochondrial protein *translocation* were repressed to a similar degree as mitochondrial ribosomal genes (Figure 5 and Table 1). Specifically, most translocation genes were repressed between 3.5-fold and 4.5-fold (mean = 3.7-fold) in day 1 adults and between 5-fold and 12-fold (mean = 7.1-fold) in day 3 adults (Table 1). This level of repression almost exactly matches the level of repression of mitochondrial ribosomal protein genes, which were repressed by an average of 3.8-fold in day 1 adults and 6.9-fold in day 3 adults (Table 1). Therefore, it appears that the mitochondrial translocase genes are co-regulated with genes involved in mitochondrial protein translation, suggesting that that mitochondrial protein *translocation* is functionally linked to mitochondrial protein *translation* in some fashion that requires them to be transcriptionally co-regulated.

### Protein Translation Genes are Not Repressed in *fem-3(q20)* Adults

It has been reported that loss of function mutations in several different types of translation genes leads to robust hypoxia resistance (Itani, Zhong et al. 2021). If reduced expression of translation genes was responsible for the primary hypoxia resistance mechanism observed in both *glp-1(e2141)* and *fem-3(q20)* mutants, then we would expect that translation genes would also be repressed in *fem-3(q20)* mutants during the L4 to adult transition. For this reason, we used RNAseq to identify changes in gene expression that occur in *fem-3(q20)* mutants between L4 and day 1 of adulthood. In contrast to our simple hypothesis, we did not find that ribosomal protein genes were significantly repressed (>2-fold) in *fem-3(q20)* adult animals (Supplementary Table 1 and Table 2). In fact, plotting the distribution of both ribosomal and mitochondrial ribosomal genes by repression shows that the distribution of ribosomal genes is indistinguishable from the distribution of total genes (Fig. 4C). In addition to the ribosomal genes, all other translation factors were also not repressed in *fem-3(q20)* mutants, except for the *let-7* microRNA (Fig 5A and 5b and Supplemental Figure 1). Interestingly, however, although the *let-7* microRNA was repressed in *fem-3(q20)* mutants, this did not lead to repression of *fib-1*, suggesting this regulatory pathway is not functioning in the *fem-3(q20)* background. Finally, we also found that genes involved in mitochondria protein translocation were not significantly repressed in *fem-3(q20)* animals upon the L4 to adult transition. These results strongly suggest that reduced expression of translation genes is not responsible for the primary hypoxia resistance observed in both *glp-1(e2141)* and *fem-3(q20)* adults upon the L4 to day 1 adult transition.

### Mutation of Protein Translation Factors Suppresses Hypoxia “Super Resistance”

The results of our *fem-3(q20)* RNAseq imply that reduced expression of translation genes is not responsible for the primary hypoxia resistance observed in both *glp-1(e2141)* and *fem-3(q20)* mutants. However, because genes involved in protein translation are repressed in *glp-1(e2141)* mutants in an aging dependent fashion, corresponding with hypoxia super resistance, we suspected that reduced protein translation may be the mechanism of hypoxia super resistance in germline deficient animals.

In a previous study it was found that mutation of two genes, *ncl-1* and *lrp-1*, was able to suppress the hypoxia resistance of multiple protein translation mutants. Specifically, mutation of both of these genes together led to an increase in the abundance of ribosomal subunits as determined by proteomic analysis (Itani, Zhong et al. 2021). To determine if *ncl-1* and *larp-1* mutations can suppress the hypoxia resistance of germline mutants, we generated the following triple mutant strains: *larp-1(q783);ncl-1(gc53);glp-1(e2141)* and *larp-1(q783);ncl-1(gc53);fem-3(q20)*. These strains were tested for hypoxia resistance as L4 larvae, day 1 adults, and day 2 adults and compared to *glp-1(e2141)* and *fem-3(q20)* single mutants. As expected, we found that dual mutation of *ncl-1* and *larp-1* did not suppress the day 1 primary hypoxia resistance of *fem-3(q20)* or *glp-1(e2141)* adults (Fig 6A and 6B). However, mutation of *ncl-1* and *larp-1* together did completely suppress the aging dependent increase in hypoxia resistance observed in *glp-1(e2141)* mutants, confirming that this aging dependent hypoxia super-resistance is likely due to reduced expression of genes involved in protein translation.

**Figure 6.**
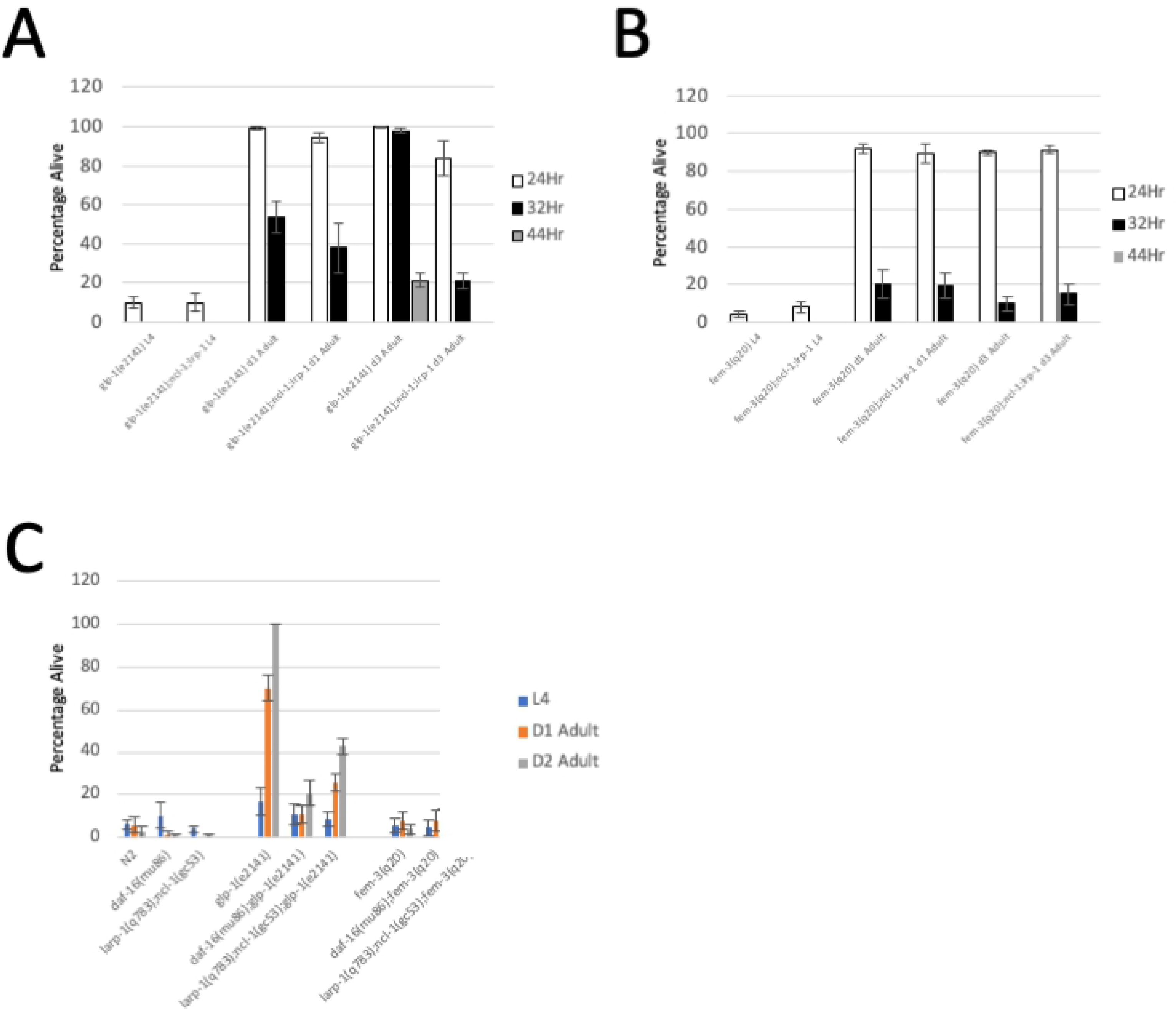
Suppressors of *glp-1(e2141)* Super Resistance. (A) Percentage of *glp-1* mutant worms surviving 24, 32, and 44 hours of hypoxia. Mutation of *ncl-1* and l*arp-1* together does not impact the day 1 adult hypoxia resistance of *glp-1(e2141)* mutants, but completely represses the aging dependent super resistance of day 2 *glp-1(e2141)* mutants. (B) Percentage of worms surviving 24, 32, and 44 hours of hypoxia. *Fem-3(q20)* mutants become more resistant to hypoxia on day 1 of adulthood, but do not become super resistant by day 2 of adulthood. Mutation of *ncl-1* and *larp-1* together does not suppress the hypoxia resistance of day 1 or day 3 *fem-3(q20)* animals. (C) Percentage of worms surviving heat stress (10.5 hours at 35°C). *glp-1(e2141)* day 1 adults survive 10.5 hours of heat stress, whereas L4 worms do not. Mutation of *daf-16* suppresses the heat resistance of *glp-1(e2141)* worms. Neither *fem-3(q20)* adults or L4 larvae survive 10.5 hours of heat stress.

### *Glp-1* Mutants are Resistant to Heat Stress as Adults but not L4 larvae

Our results demonstrate that the hypoxia resistant state of *glp-1(e2141)* doesn’t manifest until adulthood and becomes stronger as *glp-1(e2141)* mutants age. Thus, *glp-1(e2141)* L4 larvae undergo physiological changes between L4 and adulthood that confer strong resistance to hypoxic injury. This raises the possibility that the physiological properties that make *glp-1(e2141)* mutants long lived may also not occur until adulthood. This question is not easily addressed however, because it is impossible to comparatively age an L4 larva and adult. Resistance to heat stress is a marker that often correlates with longevity. Therefore, we tested larval and adult worms to determine if *glp-1(e2141)* mutants were also resistant to heat stress in a developmentally dependent manner. Worms grown at room temperature to the appropriate stage were placed at 35°C for 10.5 hours and rescued by returning to 20°C. Surviving worms were recorded 24 hours later. We found that wild type worms, *daf-16* mutants, and *larp-1;ncl-1* mutants displayed little resistance to heat stress as L4 larvae, day 1 adults or day 2 adults (Fig 6C). In contrast, *glp-1(e2141)* mutants were only mildly resistant to heat stress as L4 larvae, but became more resistant to heat stress when reaching day 1 of adulthood. The heat resistance of *glp-1(e2141)* became even stronger when reaching day 2 of adulthood. Like hypoxia resistance, the aging dependent increase in resistance to heat stress was almost fully suppressed by mutation of *daf-16* and partially suppressed by mutation of *ncl-1* and *larp-1* together (Fig 6C). In contrast, *fem-3(q20)* adults, while resistant to hypoxic injury, displayed little to no resistance to heat stress. Taken together, our results demonstrate that the primary hypoxia resistance mechanism observed in both fem-3(q20) and glp-1(e2141) mutants does not correlate with resistance to heat stress, however the *daf-16* dependent hypoxia super-resistance observed in germline deficient animals does correlate well with resistance to heat stress and, therefore, may also be related to longevity.

## DISCUSSION

### Hypoxia Resistance Correlates with Reproductive Output

Sterile mutations are sufficient to promote hypoxia resistance in *C. elegans*, as long as they prevent oogenesis or an upstream step in germline development (Mendenhall, LeBlanc et al. 2009). For this study, we set out to further establish the mechanisms by which inhibition of reproduction brings about hypoxia resistance. Using FUDR to induce partial reductions in fertility we demonstrated that total sterility is not required for hypoxia resistance. In fact, hypoxia resistance is proportional to reproductive output. The connection between fertility and hypoxia resistance is an important one because previous mutagenic and RNAi screens in *C. elegans* have identified over 200 genes that, when suppressed, lead to hypoxia resistance (Mabon, Mao et al. 2009). It was reported that inhibition of many of these genes also leads to a partial reduction in progeny production (Mabon, Mao et al. 2009). Because reproduction is an energy intensive process, the majority of a worm’s metabolic output is devoted to the production of offspring. It therefore follows that many genes that facilitate hypoxia resistance may do so simply because they restrict the amount of energy and resources available for reproduction and thereby reduce the amount of oogenesis. Our results demonstrate that this possibility should always be considered when investigating how genes influence hypoxic injury.

### A Hypoxia Sensitive to Hypoxia Resistant Transformation in Sterile Mutants

Since mutations that prevent oogenesis cause robust resistance to hypoxic injury, it is reasonable to postulate that it is the mechanism of oogenesis itself that renders worms more sensitive to hypoxia exposure. This hypothesis is not true, however, as we found that early L4 larvae that have not yet started oogenesis and WT males are just as sensitive to hypoxia as WT hermaphrodites. Furthermore, we found that *glp-1(e2141)* and *fem-3(q20)* L4 larvae, which also do not possess oocytes, are not resistant to hypoxic injury. Taken together, these data imply that it is the inhibition of oogenesis in otherwise oogenic animals that leads to hypoxia resistance, not the absence of oogenesis *per se*.

Perhaps the most interesting result of our study is the fact that the robust hypoxia resistance of sterile mutants is only observed in adult animals. In just a 12-hour period, both *glp-1(e2141)* and *fem-3(q20)* mutants transform from fully hypoxia sensitive L4 larvae to highly resistant day 1 adults. These findings imply there are considerable physiological differences between L4 larvae and day 1 adults, even in *glp-1(e2141)* mutants, which do not have an active germline. This result could be viewed as surprising, especially for *glp-1(e2141)* mutants, because the most reasonable model for why WT worms are sensitive to hypoxia and *glp-1(e2141)* mutants are not, is the presence of a germline in WT animals. In this hypothesis, it is the metabolically active germline that sends signals that make animals sensitive to hypoxia. This scenario is not true, however, as our results demonstrate that the germline itself is not required for making animals sensitive to hypoxia. Instead, we must conclude that there are developmental programs operating in somatic tissues that render animals sensitive or resistant to hypoxia. These metabolic programs may very well exist to support germline development if it were present or, alternatively, the hypoxia resistant metabolic state of sterile adults may occur as a response to the absence of the germline and/or oogenesis. In either case, these metabolic programs operate regardless of whether or not the germline is present. This is presumably true for partial sterility as well, in that partial sterility is accompanied by changes in the somatic tissues that render animals partially resistant to hypoxic injury.

### A Second Mechanism of Hypoxia Resistance Operates in Germline Ablated Animals

We also found that *glp-1(e2141)* mutants, but not *fem-3(q20)* mutants, become even more resistant to hypoxia as they age to day 2 of adulthood. In fact, over a third of *glp-1(e2141)* day 2 adults survive as much as 44 hours of hypoxia at 26.5°C. This is remarkably strong hypoxia resistance compared to WT animals, of which less than 10% survive only 20 hours of hypoxia. For clarity, we have termed this aging dependent increase in hypoxia resistance “super-resistance”. There are two lines of evidence that the super-resistance of aging *glp-1(e2141)* mutants is a second mechanism of hypoxia resistance that is additive with the primary hypoxia resistance mechanism that operates in both *fem-3(q20)* and *glp-1(e2141)* mutants. First, the primary hypoxia resistance mechanism occurs in both *glp-1(e2141)* and *fem-3(q20)* mutants, whereas the aging dependent super-resistance only operates in *glp-1(e2141)* mutants. Second, the primary hypoxia resistance mechanism of both sterile mutants does not require *daf-16*, whereas the super-resistance of *glp-1(e2141)* mutants is completely dependent on *daf-16*. Like the primary hypoxia resistance mechanism, the super resistant metabolic state must reflect a transformation in somatic tissues that occurs in the hours after *glp-1(e2141)* worms reach adulthood. Therefore, again, it is not the germline itself that determines hypoxia sensitivity, but instead it is metabolic programs that operate in the somatic tissues in response to an active or inactive germline.

### Both Hypoxia Resistance Mechanisms Prevent Mitochondrial Protein Aggregation

Protein folding errors and protein aggregation may be a key factor in hypoxic injury. It has been established that hypoxia causes protein aggregation in the mitochondria (Kaufman and Crowder 2015, Kaufman, Wu et al. 2017). Furthermore, inducing the mitochondrial unfolding response is protective against hypoxic injury. Although mitochondrial protein aggregation correlates well with hypoxia sensitivity, it is not yet known if this phenomenon is responsible for cellular toxicity. Many factors have been identified that protect against hypoxic injury, but it is not yet known if these factors prevent aggregation in the mitochondria. In this study, we found that both *fem-3(q20)* and *glp-1(e2141)* mutations prevented protein aggregation in the mitochondria. Furthermore, day 2 *glp-1(e2141)* worms were more effective at preventing aggregation than day 1 *glp-1(e2141)* worms. Therefore, it appears that both mechanisms of hypoxia protection in sterile mutants are effective at preventing mitochondrial protein aggregation. While it makes sense that reduced protein translation could protect against protein aggregation, it is not yet clear how the primary hypoxia resistance mechanisms might prevent protein misfolding in the mitochondria.

### Aging Dependent Co-Regulation of Protein Translation Genes in *glp-1(e2141)* Mutants

The fact that *glp-1(e2141)* mutants transform from hypoxia sensitive L4 larvae to hypoxia resistant day 1 adults and then to super-resistant day 2 adults provided a powerful background for identifying changes in gene expression associated with hypoxia resistance. Indeed, gene expression could be compared in the same germline deficient background at different time points. Our RNAseq studies were designed to identify gene expression changes that occur in *glp-1(e2141)* mutants between the hypoxia sensitive L4 stage, the hypoxia resistant day 1 of adulthood, and the super-resistant day 3 of adulthood. What stood out about our gene expression studies was the widespread and precise transcriptional repression of protein translation genes during the development and aging of *glp-1(e2141)* mutants. Remarkably, all 76 annotated cytosolic ribosomal protein genes (RPGs) were repressed by more than 2-fold in *glp-1(e2141)* upon the L4 to day 1 adult transition, with all being repressed to nearly the same degree (between 2.5-fold and 3-fold). Furthermore, all 64 annotated mitochondrial ribosomal genes (mitoRPGs) were repressed in day 1 adults, with nearly all repressed in a tight range (3.5-fold to 4-fold) that is more robust repression than the RPGs. These data show that cytosolic RPGs are regulated together as one group of genes (mean = 2.7-fold) and the mitochondrial RPGs (mitoRPGs) are regulated together as a second group of genes (mean = 3.8-fold repression). Both groups are repressed even more robustly in day 3 adults, with a mean of 4.0-fold for the RPGs and a mean of 6.7-fold for the mitoRPGs.

Ribosomes are very large, complex and abundant molecules that consist of multiple RNAs and dozens of distinct polypeptides. For this reason, it is generally assumed that the genes that encode ribosomal subunit proteins must be regulated together so that precise stoichiometries are achieved and ribosomes are then able to fold and assemble properly (Warner and McIntosh 2009, Sleumer, Wei et al. 2012). Incorrect stoichiometry and ribosomal mis-assembly could be consequential especially given how abundant ribosomes are in the cytosol and mitochondria. Our results here confirm that the ribosomal protein genes (RPGs) are indeed regulated together, being repressed in germline ablated animals as they transition from L4 to day 1 adults and further repressed as they age to day 3 of adulthood. We also found that the mitochondrial RPGs (mitoRPGs) are repressed under the same scenarios, arguing that there is coordination between cytosolic and mitochondrial protein translation. However, the difference in repression intensity between the RPGs and mitoRPGs suggest that the mitoRPGs are repressed via a different, more powerful, mechanism.

Cytosolic and mitochondrial translation may be precisely coordinated for good reason. Defects in the assembly and maintenance of the mitochondrial oxidative phosphorylation complex 1 lead to a range of health maladies, including diabetes, cancer, and neurodegenerative diseases (Dai and Lu 2008). Because the components of OXPHOS are translated in both the cytosol and mitochondria, it is likely that both cytosolic and mitochondrial protein synthesis is coordinated so that mitochondrial-encoded OXPHOS subunits assemble in the proper stoichiometric ratios with their nuclear counterparts.

There are two scenarios by which RPGs and mitoRPGs could be tightly regulated. First, all ribosomal protein genes could share the same transcriptional elements and transcription factors. Alternatively, ribosomal protein genes may contain a diversity of promoter elements which nevertheless result in a similar level of repression in germline ablated animals. Computational studies in *C. elegans* have identified nine distinct sequence motifs overrepresented in the promoters of RPGs (study did not focus on mitoRPGs) (Sleumer, Mah et al. 2010, Sleumer, Wei et al. 2012). However, these motifs were not found in every RPG and different combinations of these nine motifs were present in the promoters of different RPGs. Therefore, it appears that either a variety of upstream elements lead to a similar level of ribosomal gene repression in *glp-1(e2141)* mutants, or there are common regulatory sites in all RPGs that remain to be determined. Less is known about the regulation of mitoRPGs in *C. elegans*, but mitoRPGs clearly possess promoter sequences that bring about more robust repression in *glp-1(e2141)* adults than do the RPG promoters.

Our study has demonstrated that it is not just the ribosomal protein genes that are co-regulated, numerous other factors involved in cytosolic and mitochondrial protein synthesis are also repressed to a comparable degree as RPGs and mitoRPGs in *glp-1(e2141)* mutants. These factors include all 33 of the cytosolic and mitochondrial tRNA synthetases, various other ribosomal factors, nucleolar genes, as well as genes encoding mitochondrial translation initiation and elongation factors. Therefore, it appears that an entire protein translation apparatus, including both cytosolic and mitochondrial, is repressed in *glp-1(e2141)* animal as they transition from L4 larvae to day 1 adults and further repressed as they age to day 3 of adulthood.

In addition to the co-regulation of genes involved in protein translation, we also found that all 13 mitochondrial protein translocases were repressed in germline ablated animals to precisely the same degree as mitochondrial RPGs. Such strict co-regulation argues that it is essential that the stoichiometry of the mitochondrial protein translocation complex be coordinated with stoichiometry of the mitochondrial protein translation machinery. Therefore, future studies examining the co-regulation of genes involved in translation should focus on finding regulators of all protein translation genes as well as genes involved in mitochondrial translocation. Importantly, the regulation of protein translation and translocation genes occurs independently of germline signals, as their repression occurs in animals with no germline.

### Regulation of a Nucleolar Pathway in *glp-1(e2141)* Mutants

Interestingly, multiple components of a nucleolar regulatory pathway are also developmentally regulated in *glp-1(e2141)* mutants. The *let-7* microRNA is repressed by more than 100-fold in *glp-1(e2141*) adults. *Let-7* participates in nucleolar physiology by repressing the post transcriptional expression of the *ncl-1* ribosomal gene, which in turn represses the expression of *fib-1*, a gene involved in nucleolar size and function (Gruber, Ng et al. 2009, Yi, Ma et al. 2015) Repression of *let-7* would be expected to increase post transcriptional expression of the *ncl-1* gene which would in turn lead to repression of *fib-1* expression. Importantly, consistent with this model, *fib-1* gene expression is strongly repressed in *glp-1(e2141)* adults. Therefore, *let-7* functions upstream of *ncl-1* and *fib-1* to regulate nucleolar physiology in *glp-1(e2141)* mutants. It will be interesting to determine if *let-7* is an upstream regulator of all of the other translation factors in *glp-1(e2141)* adults. Importantly, *let-7* is also repressed in *fem-3(q20)* day 1 adults, yet *fib-1* and other protein translation genes are not significantly repressed in *fem-3(q20)* animals, consequently, the *let-7-ncl-1-fib-1* regulatory pathway may be modified in *fem-3(q20)* animals, such that repression of *let-7* does not lead to repression of *fib-1* and other translational proteins in this background.

### Reduced Protein Translation is the Mechanism of Super-Resistance and Longevity

It is established that mutations that reduce protein translation lead to hypoxia resistance (Itani, Zhong et al. 2021). For this reason, reduced expression of genes involved in protein translation was a good candidate for the mechanism of hypoxia resistance in germline mutants. Because we did not observe repression of protein translation genes in *fem-3(q20)* mutants, however, we must conclude that reduced expression of protein translation genes is not the mechanism for the primary hypoxia resistance observed in both *glp-1(e2141)* and *fem-3(q20)* mutants. Still, we did find that translation genes are repressed in *glp-1(e2141)* adults and further repressed as they age, therefore reduced expression of genes involved in protein translation was a candidate for the mechanism of super-resistance observed only in aging *glp-1(e2141)* mutants. Consistent with this hypothesis, we found that mutation of *ncl-1* and *larp-1* together, two repressors of protein translation, was able to suppress the hypoxia super-resistance of *glp-1(e2141*) mutants. In contrast, mutation of *ncl-1* and *larp-1* did not suppress the primary hypoxia resistance of day 1 *glp-1(e2141)* and *fem-3(q20)* adults. Thus, our experiments confirmed that reduced protein translation is the mechanism of super-resistance observed in aging *glp-1(e2141)* animals.

Reduced protein translation has also been linked to the longevity of *glp-1(e2141)* mutants (Hansen, Taubert et al. 2007). It was found that the nucleoli of different long-lived strains, including *glp-1(e2141),* are smaller than in WT worms (Tiku, Jain et al. 2017). Protein analysis determined that there were fewer ribosomal proteins present in *glp-1(e2141)* adults than there were in WT worms. Furthermore, mutation of *ncl-1* was able to suppress the longevity of *glp-1(e2141)* mutants (Tiku, Jain et al. 2017). Taken together with our results, it appears that the mechanism of hypoxia super-resistance is related, in some fashion, to the mechanism of longevity observed in *glp-1(e2141)* animals. This is further established by the fact that *daf-16* is able to suppress both longevity and super-resistance of *glp-1(e2141)* animals.

An interesting implication of our study is that the properties that enable *glp-1(e2141)* animals to live long may not be instituted in *glp-1(e2141)* worms until they reach adulthood. This hypothesis cannot be tested directly, however, because it is impossible to observe the aging of an L4 larvae without it first transforming into an adult. As a proxy to longevity, we did test the ability of larval and adult *glp-1(e2141)* animals to resist heat stress. We found that *glp-1(e2141)* are not resistant to heat stress as L4 larvae, but become resistant to heat stress as they reach adulthood and become more resistant as they age to day 2 of adulthood. These results correlate with hypoxia super-resistance. Therefore, if heat stress does indeed correlate with longevity, this would imply that only adult *glp-1(e2141)* mutants contain the physiological properties that enable them to live longer lifespans. If this true, it would establish the L4/adult transition as an excellent model for identifying the properties that enable worms to live extended lifespans.

### The Mechanism of Primary Hypoxia Resistance is Still Unknown

Our study demonstrates that there is also a profound transformation in both *glp-1(e2141)* and *fem-3(q20)* mutants between the L4 larval stage and adulthood, such that L4 larvae are fully sensitive to hypoxia injury whereas day 1 adults are highly resistant. This transition occurs in both *glp-1(e2141)* and *fem-3(q20)* adults. Since *fem-3(q20)* animals are not long lived, this second mechanism of hypoxia resistance is not a mechanism of longevity. We still do not know what the mechanism of this primary hypoxia resistance is. Unfortunately, our RNAseq analysis did not identify any specific group of genes that were induced or repressed between the L4 larval stages and day 1 of adulthood in both *glp-1(e2141)* and *fem-3(q20)* mutants. Therefore, we do not yet have a clear picture of what might be going on in the L4 to adult transition that could explain the hypoxia resistance observed in both *glp-1(e2141)* and *fem-3(q20)* adults. In a previous study, it was found that RNAi of AMP kinase subunits could suppress the *daf-16* independent hypoxia resistance of *glp-1(e2141)* mutants, implicating carbohydrate metabolism as a potential mechanism (LaRue and Padilla 2011). It remains to be determined if other sterile mutants utilize this mechanism for their primary hypoxia resistance.

### The L4/Adult Transition as a System for Studying the Unique Properties of Germline Mutants

In this study, we have determined that the hypoxia resistance of germline deficient mutants is not a general property, but instead hypoxia resistance is “switched on” only in adult animals. This observation demonstrates that the signals that determine hypoxia sensitivity are derived from somatic tissues and act in developmental fashion. This study therefore pinpoints the L4/adult transition of germline deficient mutants as an excellent system to study the unique physiological properties of germline mutants. Adult germline mutants can simply be compared to L4 larvae to identify physiological changes that determine hypoxia sensitivity. This is a much cleaner system than comparing germline deficient mutants to WT animals. In this study, we used such a comparison to identify reduced expression of genes involved in protein translation as one of the mechanisms of hypoxia resistance in germline deficient animals. Our data also suggest that this same phenomenon may be true for the long-lived properties of germline deficient animals. Future studies should focus on the L4/adult transition to identify physiological changes associated with hypoxia resistance, heat stress resistance, and longevity.

## SUPPLEMENTARY DATA

Figure S1. Repression of Protein Translation and Nucleolar Genes in Germline Mutants. Fold repression of tRNA synthetases (top) ribosomal factors (middle) and genes involved in nucleolar biology (bottom). All three of these gene classes show repression between L4 and day 1 of adulthood in glp-1(e2141) mutants, but no increase in repression between day 1 and day 3 of adulthood in glp-1(e2141) mutants. No significant repression is observed in fem-3(q20) mutants between L4 and day 1 of adulthood.

## SUPPLEMENTARY TABLES

**Table S1. Repression of ribosomal protein genes (RPGs) in *glp-1(e2141)* and *fem-3(q20)* mutants.**

**Table S2. Repression of mitochondrial ribosomal protein genes (mRPGs) in *glp-1(e2141)* and *fem-3(q20)* mutants.**

## METHODS

### *C. elegans* strains and maintenance

**N2-Bristol**, **AGD1032:**glp-1(e2141), **CF1880:**daf-16(mu86);glp-1(e2141), **HGA8006:**glp-1(e2141);fat-5(tm420), **HGA8007:**glp-1(e2141);fat-6(tm331), **HGA8008:**glp-1(e2141);fat-7(wa36), **CB3844:**fem-3(e2006), **JK816:**fem-3(q20), and **SS104:**glp-4(bn2) were obtained from the *C. elegans* Genetics Center at the University of Minnesota. **MC893:**larp-1(q783);ncl-1(gc53) was obtained as described. The following compound mutant strains were constructed using standard genetic techniques: **MC952:**larp-1(q783);ncl-1(gc53);glp-1(e2141), **MC942:**larp-1(q783);ncl-1(gc53);fem-3(q20), and **MC965:**daf-16(mu86);fem-3(q20).

Animals were maintained at 20°C on nematode growth media (NGM) plates seeded with Escherichia coli OP50. The N2 (Bristol) strain was the standard wild type strain from the *C. elegans* Genetics Center (CGC, University of Minnesota). Compound mutants were constructed using standard genetic techniques. Genotypes were confirmed by PCR amplification or by PCR followed by restriction digest.

### Hypoxia Assays

For hypoxia assays, worms were synchronized by allowing 5 hermaphrodites to lay eggs for 4 hours. Adult hermaphrodites were removed from plates and synchronized progeny were grown at 25°C until ready for assay. Worms were assayed as either L4 larvae, day 1 adults or day 2 adults. Day 1 adults were assayed ∼12-24 hours after the L4/adult transition, day 2 adults were assayed 36-48 hours after the L4/adult transition. Synchronized young adult worms were subjected to hypoxia as described previously except that hypoxic incubation temperature was 26.5°C (Mao and Crowder 2010). Briefly, each plate of worms was washed into one 1.5 ml tube with 1 ml of M9 buffer (22 mM KH2PO4, 22 mM Na2HPO4, 85 mM NaCl, 1 mM MgSO4). Worms were allowed to settle by gravity, and 900 μl of M9 buffer was removed. The tubes were then placed in the anaerobic chamber (Forma Scientific) at 26.5°C for the indicated incubation times. Oxygen tension was always ≤ 0.3%. Following the hypoxic insult, worms were placed using glass Pasteur pipettes onto NGM plates spotted with OP50 bacteria and recovered at 20°C for 24 hours. Worms that moved or responded to being prodded with a pick were scored as alive.

### FUDR Assays

Worms were synchronized by allowing 5 hermaphrodites to lay eggs for 4 hours. When progeny reached the L4 stage of development, the appropriate concentration of FUDR was added to the growth plates. Worms were then grown to the appropriate stage, placed in hypoxia chamber and assayed as described above.

### Analysis of Protein Aggregation

Wild type and mutant worms were synchronized as described above and grown to day 1 or day 2 of adulthood. Worms were then exposed to hypoxia at 26.5°C. Immediately after hypoxia exposure, worms were mounted for microscopy. Confocal microscopy was performed using a Leica TCS SP8 equipped with a HyD detector. Paralysis was produced by mounting worms in a solution of 50 mM levamisole (Sigma-Aldrich Corp., St. Louis, MO, USA) in M9 prior to imaging. Images were acquired from at least ten randomly selected worms at 1024 × 1024 resolution using a 63 × objective with 8 × zoom producing a 23.07 × 23.07 *μ*m image (defined as one high power field, HPF). All images were acquired as a ten slice Z-stack with scan speed of 800-1800 Hz and flattened as a maximum intensity projection prior to analysis. Analysis of images was performed by an observer blinded to condition. For assessment of total fluorescence images were acquired on a Zeiss Axioskop2 microscope with fluorescence intensity quantified using NIH ImageJ.

### Heat Stress Assays

Worms were synchronized by allowing 5 adults lay eggs for 4 hours. Synchronized progeny were grown to the L4, day 1, or day 2 stage of adulthood on NGM plates with OP50 and then were placed in a 35°C incubator for 10.5 hours. Worms were then returned to 20°C and scored for survival 24 hours later.

### RNAseq

*C. elegans* strains *glp-1(e2141)* and *fem-3(q20)* were grown at 20°C on high-growth plates seeded with OP50 bacteria and maintained as described (Van Gilst, Hadjivassiliou et al. 2005). Gravid adults from 10 10-cm plates were bleached, and embryos were dispersed onto 15-cm nematode growth media (NGM)-lite plates seeded with OP50. ∼10,000 Worms at the L4 stage, day 1 stage and day 2 stage were harvested, washed twice with M9, and frozen in liquid nitrogen. For RNA preparation, worms were thawed at 65°C for 10 min, and RNA was isolated using the Tri-Reagent Kit (Molecular Research Center, Cincinnati, Ohio, United States). RNAseq was performed by NovoGene Corporation, Sacramento, California. Read count files obtained from NovoGene were normalized to a panel of genes previously shown to be stably expressed under a variety of conditions (Van Gilst, Hadjivassiliou et al. 2005, Van Gilst, Hadjivassiliou et al. 2005, Pathare, Lin et al. 2012). 4 data sets were collected for *glp-1(e2141)* L4, day 1 adult, and day 2 adult. 3 data sets were collected for *fem-3(q20)* L4 and day 1 adults. The genes used for normalization were as follows: *lbp-3, acdh-7, pod-2, fasn-1, nhr-80, pfk-1, pdk-2,* Y110A7A.6, *gpd-3, gpd-2, aco-2, and F48E8.3*. The following formula was used for normalization…

> 1 / ((REF_lbp-3_/DATA_lbp-3_ + REF_*acdh-7*_/DATA_*acdh-7*_ + REF_*pod-2*_/DATA_*pod-2*_ +
>
> REF_*fasn-1*_/DATA_*fasn-1*_ + REF_*nhr-80*_/DATA_*nhr-80*_ + REF_*pfk-1*_/DATA_*pfk-1*_ + REF_*pck-*_
>
> _*2*_/DATA_*pck-2*_ + REF_*Y110A7A.6*_/DATA_*Y110A7A.6*_ + REF_*gpd-3*_/DATA*gpd-3* + REF_*gpd-*_
>
> _*2*_/DATA_*gpd-2*_ + REF_*aco-2*_/DATA*_aco-2_* + REF*_F48E8.3_*/DATA*_F48E8.3_*) / 12) = AF

Where REF is the read count for the reference data set, and DATA is the read count for the data set being normalized.

> Final read count was determined by the following formula
>
> CRC*_i_* = NF*RRC*_i_*

Where CRC is the corrected read count for gene *i*. NF is equal to the normalization factor and RRC*_i_* is the raw read count for gene *i*.

> Fold repression was determined by the following formula:
>
> FR(L4->D1)_i_ = (*glp-1* L4 CRC)_i_/(*glp-1* D1 CRC)_i_

Where FR(L4->D1) is fold repression of gene i between L4 and day 1 of adulthood. (*glp-1* L4 CRC)_i_ is the corrected read count for gene i from *glp-1(ts)* L4 worms, and (*glp-1* D1 CRC)_i_ is the corrected read count for gene i from *glp-1(ts)* D1 adults. For day 3 repression, REP(L4->D3, (*glp-1* D1 CRC)_i_ is substituted with (*glp-1* D3 CRC)_i_. The same methodology was used to determine fold repression between L4 and day 1 of adulthood in *fem-3(q20)* worms. Fold activation was calculated by the following formula:

> FA(L4->D1) = 1/FR(L4->D1).

Where FA(L4->D1) is the fold activation between L4 and day 1 of adulthood.

## Notes

### Competing Interest Statement

The authors have declared no competing interest.

